# When to use Quantile Normalization?

**DOI:** 10.1101/012203

**Authors:** Stephanie C. Hicks, Rafael A. Irizarry

## Abstract

Normalization and preprocessing are essential steps for the analysis of high-throughput data including next-generation sequencing and microarrays. Multi-sample global normalization methods, such as quantile normalization, have been successfully used to remove technical variation from noisy data. These methods rely on the assumption that observed global changes across samples are due to unwanted technical variability. Transforming the data to remove these differences has the potential to remove interesting biologically driven global variation and therefore may not be appropriate depending on the type and source of variation. Currently, it is up to the subject matter experts, for example biologists, to determine if the stated assumptions are appropriate or not. Here, we propose a data-driven method to test for the assumptions of global normalization methods. We demonstrate the utility of our method (*quantro*), by applying it to multiple gene expression and DNA methylation and show examples of when global normalization methods are not appropriate. We also perform a Monte Carlo simulation study to illustrate how our method generally outperforms the current approach. An R-package implementing our method is available on Bioconductor (http://www.bioconductor.org/packages/release/bioc/html/quantro.html).

## Introduction

Multi-sample normalization techniques such as quantile normalization [1, 2] have become a standard and essential part of analysis pipelines for high-throughput data. These techniques transform the original raw data to remove unwanted *technical variation*. Technical variation can cause perceived differences between samples processed on high-throughput technologies, irrespective of the biological variation. These differences are typically due to changes in experimental conditions that are hard or impossible to control [3] and confusing them with biological variability can lead to false discoveries [4, 5].

Some of the first attempts at normalizing microarray data mimicked the use of so-called house-keeping genes [6] as was done by the established gene expression measurement technology that preceded microarrays. This approach did not work well in practice [7, 8], therefore data-driven approaches were developed such as median correction [9, 10], variance-stabilizing transformation [11], locally weighted linear regression (loess) [12] and spline based methods [13]. The general idea of these approaches is to assume that observed variability in global properties are due only to technical reasons and are unrelated to the biology of interest [2, 14]. Here we refer to these as *global adjustment* methods [15]. Examples of global properties include the total number of differentially expressed genes across groups, the median gene expression across genes and the statistical distribution of gene expression values. These types of assumptions are justified in many biomedical applications, for example in gene expression studies in which only a minority of genes (or *targeted* set of genes) are expected to be differentially expressed. However, if, for example, a substantially higher percentage of genes are expected to be expressed in only one group of samples, it may not be appropriate to use global adjustment methods.

Quantile normalization was originally developed for gene expression microarrays [1, 2] but today it is applied in a wide-range of data types including genotyping arrays [16, 17], RNA-Sequencing (RNA-Seq) [18-20], DNA methylation [21], ChIP-Sequencing [22, 23] and brain imaging [24-26]. Quantile normalization is a global adjustment method that assumes the statistical distribution of each sample is the same. Normalization is achieved by forcing the observed distributions to be the same and the average distribution, obtained by taking the average of each quantile across samples, is used as the reference. This method has worked very well in practice but note that when the assumptions are not met, global changes in distribution that may be of biological interest will be wiped out and features that are not different across samples can be artificially induced [27]. A schematic of quantile normalization is provided in Figure 1.

**Figure 1:**
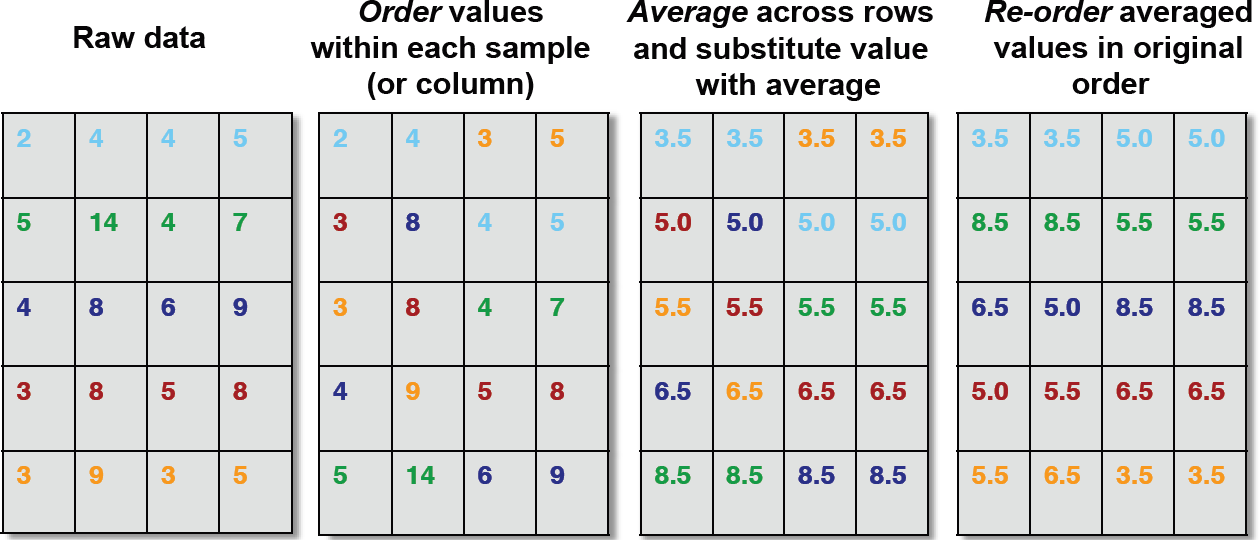
A schematic of quantile normalization. Quantile normalization is a nonlinear transformation that replaces each feature value (row) with the mean of the features across all the samples with the same rank or quantile. To quantile normalize a raw high-throughput data set with multiple samples: (1) order the feature values within each sample (2) for each feature, average across the rows (3) substitute the raw feature value with the average (4) re-order the transformed values by placing in the original order.

Previously, the burden of deciding if these assumptions hold have been left to the experimentalist. Here we propose a statistical test, referred to as *quantro*, for the assumptions of global adjustment methods, such as quantile normalization, that tests for global differences in distributions between groups of samples. Our test uses the raw unprocessed high-throughput data as input to calculate a test statistic comparing the variability of distributions within groups relative between groups. If the variability between groups is sufficiently larger than the variability within groups, then this suggests there may be global differences in distributions between groups of samples and global adjustment methods may not be appropriate. We demonstrate the advantages of our method by applying it to several gene expression and DNA methylation datasets with *targeted* and *global* changes in distributions. We define *global* changes as an abundance of differences between two or more sets of samples affecting the shape or the location shift of the distributions across groups caused by a biological or a technical source of variation and *targeted* changes as differences between sets of samples not affecting the shape or location shift of the distributions caused by a biological or a technical source of variation. We also perform a Monte Carlo simulation study to illustrate how our method generally outperforms the current approach when there are global biological differences in the distributions between groups.

## Results

### quantro: Test for global differences in distributions between groups

Consider a set of raw high-throughput data *X*_*ik*_ representing *i* ∈ (1,…, *n*_*k*_) samples in each of the *k* ∈ (1, …, *K*) groups (*n*_*T*_ total samples) from a gene expression or DNA methylation experiment. We assume *X*_*ik*_ has some common distribution 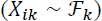 where 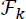 is the theoretical distribution for the *k*^*th*^ group. We define 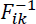 as the observed quantile distribution for the *i*^*th*^ sample in the *k*^*th*^ group. As a first step, we use an ANOVA to test if the average of the medians of the distributions are different across groups and median normalize the samples accordingly. Let 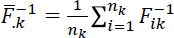 be the quantile distribution averaged across all samples in the *k*^*th*^ group and let 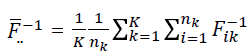 be the quantile distribution averaged across all samples and groups.

To quantify the differences between two distributions, we use Mallow’s distance [28], which is defined as the distance between two probability distributions over a region (Supplementary Eq. 1). We define the *total variance* of the distributions as the sum of squared differences between 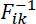 and 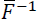 using Mallow’s distance (in the case where *p* = 2) as 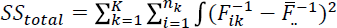. The *total variance* can be decomposed (Supplementary Eqs. 2-7) into the variance between groups (*SS*_*between*_) and the variance within groups (*SS*_*within*_):

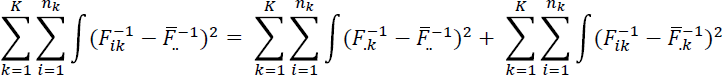

We propose using a data-driven test statistic, referred to as *F*_*quantro*_, to test for global differences in the distributions between the *K* groups. The null hypothesis is that there are no global differences in the distributions between the groups and the alternative hypothesis is that at least one group is different from the rest.

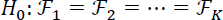

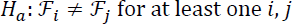

If there are no global differences in the distributions between the groups (due to technical or biological variation), we can apply a global adjustment method, such as quantile normalization, to remove any unwanted technical variation. If there are global differences in the distributions between the groups, quantile normalization may not be an appropriate normalization technique depending on the source of variation (technical or biological variation).

The *F*_*quantro*_ test statistic (Supplementary Eq. 8) is a ratio of the mean squared error between groups (*MS*_*between*_) to the mean squared error within groups (*MS*_*within*_):

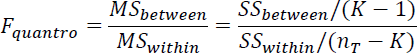

We use permutation testing to assess the statistical significance of *F*_*quantro*_ and reject the null hypothesis if the *p*-value (Supplementary Eq. 9) from the permutation test is less than some *α* significance level.

### Targeted and global changes in gene expression

We applied *quantro* to several publicly available gene expression datasets based on both microarray and RNA-Seq platforms (Supplementary Table 1) to investigate *targeted* and *global* differences in distributions across groups. We used an *α* = 0.05 significance level as the threshold to test for *global* changes in the distributions across groups. Examples of targeted changes in distributions across groups are the gene expression of samples from the Yoruba (YRI) population stratified by genotype based on an expression quantitative trait loci (eQTL) (*p* = 0.917, Figure 2a and Supplementary Fig. 1), samples from two inbred mouse strains (*p* = 0.245, Supplementary Fig. 2), samples of alveolar macrophages from nonsmokers, smokers and patients with asthma (*p* = 0.562, Figure 2b and Supplementary Fig. 3), samples of bronchial brushings from individuals with and without COPD (*p* = 0.218, Supplementary Fig. 4) and samples from two regions of the brain in patients with Parkinson’s disease (*p* = 0.264, Supplementary Fig. 5). In all of the above examples, quantile normalization is considered appropriate because no global differences in the distributions across groups were detected at the *α* = 0.05 significance level.

**Figure 2:**
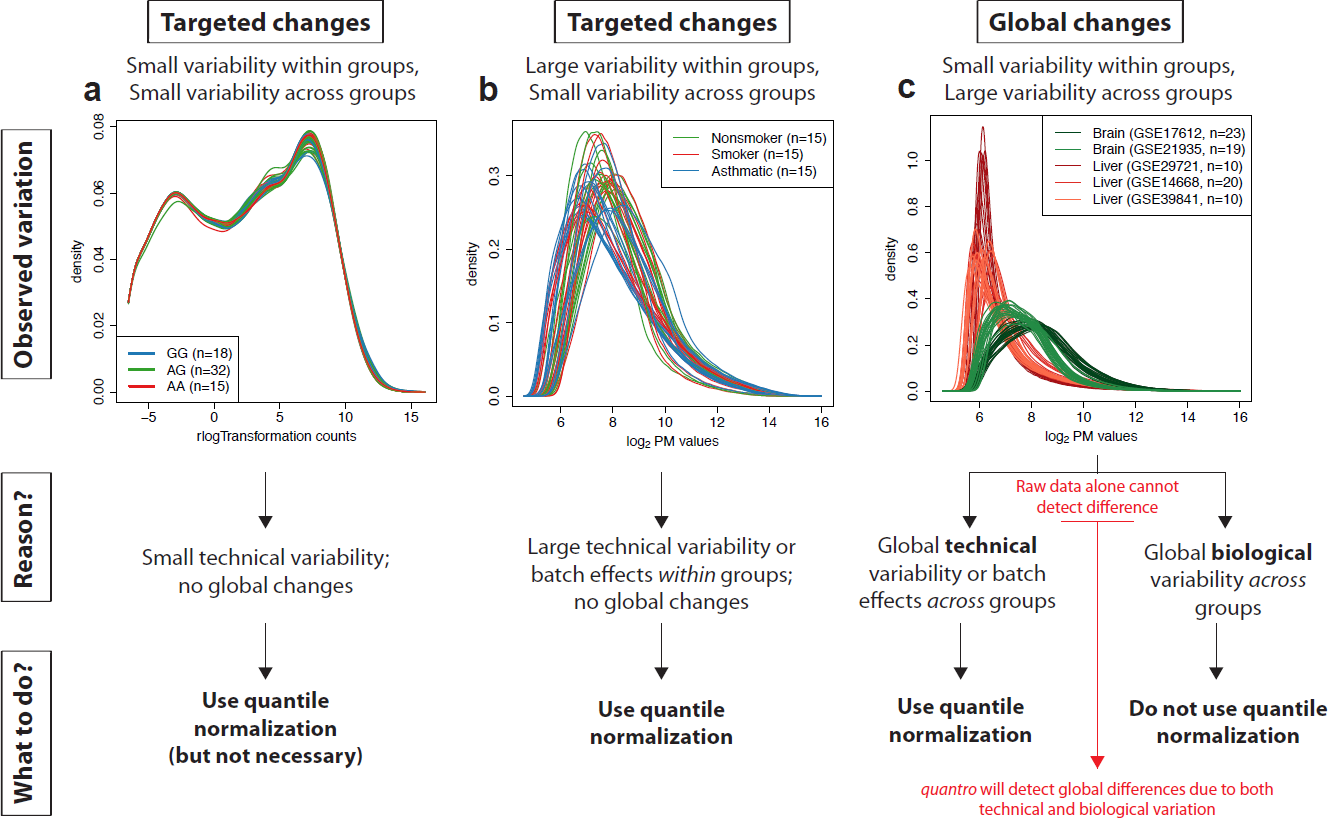
When to use quantile normalization? Examples of gene expression data with *targeted changes* and *global changes* in distributions across groups. (**a**) Transformed read counts from *n* = 65 RNA-Seq samples from the YRI population and colored by genotype based on the eQTL rs7639979: GG (blue), GA (green) and AA (red). As no global differences in distributions were detected, this suggests quantile normalization is appropriate, but not necessary as there is a low level of variation within and between groups. (**b**) Raw PM values from *n* = 45 arrays comparing the gene expression of alveolar macrophages from nonsmokers (green), smokers (red) and patients with asthma (blue). No global differences in distributions were detected which indicates quantile normalization is appropriate, as it will remove any platform-based technical variability or batch effects within groups. (**c**) Raw PM values from *n* = 82 arrays comparing brain and liver tissue samples. The samples are colored by tissue (brain [red] and liver [green]), and the shades represent different GEO IDs. The global differences in distributions detected across brain and liver tissues indicate quantile normalization is not appropriate. Global changes caused by technical variation (e.g. batch effects across groups) will also be detected by *quantro*, but raw data alone cannot detect this difference.

When comparing the gene expression of two tissues, we found striking global differences in the distributions between brain and liver tissues (*p* = 0.004, Figure 2c and Supplementary Fig. 6). We considered multiple studies from GEO to represent each tissue to prevent batch effects [29] of different studies from GEO being confounded with differences in tissues. We also compared the gene expression of normal and tumor samples. We obtained multiple studies from GEO and found global differences in the distributions between the normal and tumor samples of lung (*p* < 0.001, Figure 2D), breast (*p* < 0.001), prostate (*p* < 0.001), thyroid (*p* < 0.001), stomach (*p* < 0.001) and liver tissues (*p* = 0.044) (Supplementary Figs. 7-12). We also found global changes in the distributions of liver tissues between four groups of patients (control, healthy obese, steatosis and nash samples) from a study investigating the gene expression of NonAlcoholic Fatty Liver Disease (*p* = 0.004, Supplementary Fig. 13).

### Targeted and global changes in DNA methylation

In addition to gene expression, we considered three publicly available DNA methylation data sets. We detected no global differences in distributions of adipose tissues from patients before and after six months of exercise (*p* = 0.132, Figure 3a and Supplementary Fig. 14) and pancreatic tissues from non-diabetic and Type 2 diabetes (T2D) (*p* = 0.069, Supplementary Fig. 15). In contrast, *quantro* detected global differences in the distributions across six purified cell types from whole blood (*p* < 0.001, Figure 3b and Supplementary Fig. 16), which may be relevant for the studies estimating the cell composition of whole blood using DNA methylation [30, 31].

**Figure 3:**
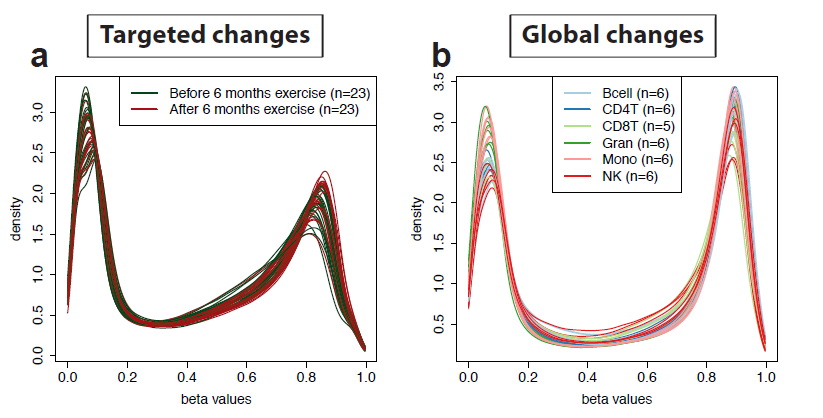
Biological variation in distributions of raw DNA methylation microarrays. (a) Example of *targeted changes* in distributions: raw beta values from *n* = 46 arrays comparing adipose tissue samples from healthy men before and after 6 months of exercise. Global changes: (b) Example of *global changes* in distributions: raw beta values from *n* = 35 arrays comparing six purified cell types from whole blood: CD14+ Monocytes (Mono), CD19+ B-cells (Bcell), CD4+ T-cells (CD4T), CD56+ NK-cells (NK), CD8+ T-cells (CD8T), and Granulocytes (Gran).

### *quantro* improves the accuracy of detecting differentially methylated CpGs

Here we evaluate the performance of using *quantro* in the context of *targeted* and *global* changes in distributions with the goal of detecting differentially methylated CpGs. We performed a Monte Carlo simulation study to compare the relative improvement of using *quantro* to the naïve approach of always using quantile normalization where *quantro* uses the *F*_*quantro*_ test statistic to decide if quantile normalization is appropriate (no normalization otherwise). For the simulation study, we simulate DNA methylation arrays with a goal of detecting differentially methylated CpGs, but note these results also translate for differential gene expression.

If there is only a minority of differentially methylated CpGs, quantile normalization reduces the bias and mean squared error (MSE) in detecting true differences between groups of samples because it removes unwanted technical variation (Supplementary Fig. 21-22). As the number of differentially methylated CpGs increases, quantile normalization will remove both the unwanted technical and interesting biological variation resulting in higher bias and MSE when detecting differential methylation. In contrast, *quantro* reduces the bias and MSE compared to using quantile normalization because the method is able to detect when there are global differences (Supplementary Fig. 21-22). Similarly, the number of false discoveries is reduced when using *quantro* when there are global differences between groups. For example, when considering a 450K DNA methylation array if there are only a small number of differentially methylated CpGs (1% of CpGs or 4,500 CpGs), *quantro* and quantile normalization are comparable in the number of false discoveries (873 and 873, respectively), but if there are global differences in the distributions between groups (10% of CpGs or 45,000 CpGs), *quantro* is able to detect those global differences and reduce the number of false discoveries compared to quantile normalization (4887 and 6583, respectively) (Supplementary Fig. 23). Using *quantro* to test if global normalization methods are appropriate gives researchers a data-driven test that yields smaller bias, MSE and less false discoveries compared to naively using quantile normalization when detecting differentially methylated CpGs.

In addition, we considered the true positive rate and false positive rate of the two normalization approaches while varying the threshold of the number of top differentially methylated CpGs selected. If there are only a small number of differentially methylated CpGs, quantile normalization and *quantro* are comparable in performance, but when the proportion of differentially methylated CpGs increases, *quantro* outperforms using quantile normalization because it can detect when there are global differences between groups (Supplementary Fig. 24). Using *quantro* as a tool to determine which type of normalization approach to employ results in higher sensitivity and specificity when detecting true differentially methylated CpGs compared to naively using quantile normalization.

## Discussion

The advent of high-throughput technologies brought the opportunity for researchers to investigate and assess biological variability at the genomic level, but it also introduced unwanted technical variability that can cause perceived differences between samples processed on high-throughput technologies, irrespective of the biological variation. These differences may be due to differences in the way the samples were processed (such as batch effects) or be due to platform-dependent technical variation. Because global changes in distributions between groups can be caused by both technical variation and biological variation, it is important to note that our test statistic *F*_*quantro*_ will detect global differences caused by both technical variation (e.g. batch effects) and biological variation. Data alone cannot determine if global changes are caused by technical variation or biological variation (Figure 2), but *quantro* offers researchers a new tool to detect when there are global changes in distributions across groups.

Here we have shown if there are global changes in the distributions across a set of groups, normalization methods with *global adjustments* may not be appropriate depending on the type and source of variation. If global adjustment methods are not appropriate, other methods such as *application-specific* methods [15] can be used. These are normalization methods where the adjustments are directly incorporated into the experiment or main analysis. Examples of these methods include the use of positive and negative control genes, the use of spike-in controls and explicitly modeling known or unknown effects of unwanted variation in a linear model (see Supplementary Section 5 for more a more detailed discussion on application-specific methods).

Previous studies have evaluated and discussed normalization methods with and without global adjustments [2, 15, 32], but the decision of which type of normalization method to use depends on the outcome of interest. For example, a recent study [27] discussed the use of normalization procedures in global gene expression analysis comparing two schematics: targeted changes in gene expression and global changes in gene expression such as transcriptional amplification [33] or transcriptional shutdown [34]. Not surprisingly, the authors show normalization methods with global adjustments are not appropriate if the total RNA is not the same across the samples. In this case, if normalization is performed at the experimental level (introducing similar amounts of RNA into the assay from the two groups with global changes), then we suggest using control genes or spike-ins controls as no differences between the distributions will be detected (Supplementary Fig. 25). However, for the great majority of studies such strategies are not available. Furthermore, if one knows *a priori* that most genes are differentially expressed then it is not clear why one would use these high-throughput technologies.

To test if normalization methods with *global adjustments* are appropriate, we developed method to test *a priori* to the data analysis for the assumptions of global normalization methods, such as quantile normalization. We have demonstrated the utility of our method by applying it to several gene expression and DNA methylation datasets revealing examples of both targeted and global changes in distributions across groups such as the global changes in distributions detected between the gene expression of brain and liver tissues. We demonstrated that *quantro* outperforms the current approach because the method is able to detect when there are global differences in distributions and therefore prevent removing potentially interesting biological variation. We have implemented our method into the *quantro* R-package providing researchers a tool to test the assumptions of global normalization methods in the analysis of their own data.

## Methods

### Data analysis

The method introduced here has been implemented into the *quantro* R-package available on Bioconductor. To test for global differences in distributions between groups of samples from high-throughput data sets, we applied *quantro* to several publicly available gene expression and DNA methylation data sets. Supplementary Table 1 contains a list of all the data sets. For this analyses, we use the *α* = 0.05 significance level as the threshold to detect *global* changes in the distributions across groups.

To compare the gene expression on microarrays of cancer samples and brain and liver tissues, we considered multiple studies from GEO [35] to represent each tissue to prevent batch effects of different studies from GEO being confounded with differences between cancer samples or between tissues. For the gene expression samples using microarrays, we extracted the raw Perfect Match (PM) values from the CEL files using the *affy* R/Bioconductor package [36]. To visualize the true biological variation in the experimentally normalized samples from Loven et al. (2012), we divided the raw PM values by the sample mean of the PM values across the spike-ins on the log_2_ scale. For the gene expression samples using RNA-Seq, we used the rlogTransformation provided in the *DESeq2* R/Bioconductor package [37] to transform the raw counts to the log_2_ scale. The RNA-Seq data was obtained from ReCount [38], which pre-processes the raw sequencing data and provides a table of raw counts for each gene. We removed all the rows with zero counts across all the samples. For the DNA methylation samples using microarrays, we used the *minfi* R/Bioconductor package [39]. We extracted the raw methylated and unmethylated signal using and computed the “beta”-values using Illumina’s default setting of the *offset* parameter equal to 100.

### Details for simulation studies

We developed an R package, referred to as *quantroSim* (Supplementary Section 3), which is available on github, to simulate gene expression and DNA methylation data, but here we just focus on DNA methylation. To simulate samples on a microarray platform technology, we use the Langmuir adsorption model [40] to model the chemical saturation in the hybridization of the probes. Each of the simulation studies considered two groups with five samples each (total of 10 samples).

With the goal of detecting differentially methylated CpGs, we compared the performance of *quantro* to the naïve approach of always using quantile normalization where *quantro* uses the *F*_*quantro*_ test statistic to decide if quantile normalization is appropriate (no normalization otherwise) (Supplementary Section 4). For the permutation testing in *quantro*, we used 100 permutations and a cutoff threshold of *α* = 0.05, unless specified otherwise. After normalization, the difference between the group means were estimated and the top differentially methylated probes were found using a *t*-test.

We assessed the relative bias (bias from *quantro* to the bias from quantile normalization) and relative mean squared error (MSE) while varying the cutoff threshold *α* from *quantro* and for a fixed threshold at *α* = 0.05. We simulated DNA methylation samples with a varying proportion of differentially methylated CpGs between the two groups and a varying level of technical variation (see Supplementary Sections 3 and 4 for more details).

To select a list of top differentially methylated probes, we adjusted the *p*-values from a *t*-test using the Benjamini & Hochberg adjustment to correct for multiple testing. The number of false discoveries was calculated using as the number of incorrectly selected probes from a given set of top differentially methylated probes. The true positive rate (TPR) was calculated as the number of correctly selected probes from the set of true differentially methylated probes. In contrast, the false positive rate (FPR) was calculated as the number of incorrectly selected probes from the set of probes that are not differentially expressed.

## Software

The R-package *quantro* implementing our method is available in Bioconductor 3.0 (http://www.bioconductor.org/packages/release/bioc/html/quantro.html) and the *quantroSim* R-package to simulate gene expression and DNA methylation data is available on Github (https://github.com/stephaniehicks/quantroSim).

## Supplementary Material

Supplementary materials are available in a single pdf. All scripts containing the code for these analyses are available on Github http://stephaniehicks.github.io/quantroPaper/

## Authorship Contribution

S.C.H and R.A.I. developed the method *quantro* and the models used in *quantroSim.* S.C.H. wrote the *quantro* and *quantroSim* R packages, analyzed the gene expression and DNA methylation data and performed the simulation studies. S.C.H. and R.A.I. wrote the manuscript.

## Competing Financial Interests

The authors declare no financial competing interests.

## Funding

S.C.H and R.A.I. were supported by NIH R01 grants GM083084 and RR021967/GM103552.

## References

1. Amaratunga, D., Cabrera, J. Analysis of Data From Viral DNA Microchips. J Amer Statist Assoc 96, 1161–1170 (2001).

2. Bolstad, B.M., Irizarry, R.A., Astrand, M., Speed, T.P. A comparison of normalization methods for high density oligonucleotide array data based on variance and bias. Bioinformatics 19, 185–193 (2003).

3. Scherer, A. Batch effects and noise in microarray experiments. John Wiley & Sons, Chichester, United Kingdom (2009).

4. Allison, D.B., Cui, X., Page, G.P., Sabripour, M. Microarray data analysis: from disarray to consolidation and consensus. Nat Rev Genet 7, 55–65 (2006).

5. Auer, P.L., Doerge, R.W. Statistical design and analysis of RNA sequencing data. Genetics 185, 405–416 (2010).

6. Butte, A.J., Dzau, V.J., Glueck, S.B. Further defining housekeeping or “maintenance,” genes focus on “Aacompendium of gene expression in normal human tissues”. Physiol Genomics 7, 95–96 (2001).

7. Eisenberg, E., Levanon, E.Y. Human housekeeping genes are compact. Trends in Genet 19, 362–365 (2003).

8. Eisenberg, E., Levanon, E.Y. Human housekeeping genes, revisited. Trends in Genet 29, 569–574 (2011).

9. Cho, R.J. et al. A genome-wide transcriptional analysis of the mitotic cell cycle. Mol Cell 2, 65–73 (1998).

10. Selinger, D.W. et al. RNA expression analysis using a 30-base pair resolution *Eschericha coli* genome array. Nat Biotechnol 18, 1262–1268 (2000).

11. Durbin, B.P., Hardin, J.S., Hawkins, D.M., Rocke, D.M. A variance-stabilizing transformation for gene-expression microarray data. Bioinformatics 18, S105–110 (2002).

12. Yang, Y.H., et al. Normalization for cDNA microarray data: a robust composite method addressing single and multiple slide systematic variation. Nucleic Acids Res 30, e15 (2002).

13. Workman, C. et al. A new non-linear normalization method for reducing variability in DNA microarray experiments. Genome Biol 3, research0048.1–research0048.16 (2002).

14. Reimers, M. Making informed choices about microarray data analysis. PLoS Comput Biol 6, e1000786 (2010).

15. Gagnon-Bartsch, J.A., Speed, T. Using control genes to correct for unwanted variation in microarray data. Biostatistics 13, 539–552 (2012).

16. Carvalho, B.S., Louis, T.A., Irizarry, R.A. Quantifying uncertainty in genotype calls. Bioinformatics 15, 242–249 (2010).

17. Scharpf, R.B., Irizarry, R.A., Ritchie, M.E., Carvalho, B., Ruczinski, I. Using the R package crimm for genotyping and copy number estimation. Journal of Statistical Software 40, 1–32 (2011).

18. Cloonan, N., et al. Stem cell transcriptome profiling via massive-scale mRNA sequencing. Nat Methods 5, 613–619 (2008).

19. Bullard, J.H., Purdom, E., Hansen, K.D., Dudoit, S. Evaluation of statistical methods for normalization and differential expression in mRNA-Seq experiments. BMC Bioinformatics 11, 94 (2010).

20. Dillies, M.A. et al. A comprehensive evaluation of normalization methods for Illumina high-throughput RNA sequencing data analysis. Brief Bioinform 14, 671–683 (2013).

21. Yousefi, P. et al. Considerations for normalization of DNA methylation data by Illumina 450K BeadChip assay in population studies. Epigenetics 8, 1–12 (2013).

22. Bilodeau, S., Kagey, M.H., Frampton, G.M., Rahl, P.B., Young, R.A. SetDB1 contributes to repression of genes encoding developmental regulators and maintenance of ES cell state. Genes & Dev 23, 2484–2489 (2009).

23. Kasowski, M. et al. Variation in transcription factor binding among humans. Science 328, 232–235 (2010).

24. Nyúl, L.G., Udupa, J.K., Zhang, X. New variants of a method of MRI scale standardization. IEEE Trans Med Imaging 19, 143–150 (2000).

25. Shah, M. et al. Evaluating intensity normalization on MRIs of human brain with multiple sclerosis. Med Image Anal 15, 267–282.

26. Shinohara, R.T. et al. Statistical normalization techniques for magnetic resonance imagine. Neuroimage Clin (inpress) (2014).

27. Loven, J. et al. Revisiting global gene expression analysis. Cell 151, 476–482 (2012).

28. Mallows, C.L. A note on asymptotic joint normality. Ann. Math. Statist. 43, 508–515 (1972).

29. Leek, J.T. et al. Tackling the widespread and critical impact of batch effects in high-throughput data. Nat Rev Genet 11, 733–739 (2010).

30. Houseman, E.A. et al. DNA methylation arrays as surrogate measures of cell mixture distributions. BMC Bioinformatics 13, 86 (2012).

31. Koestler, D.C. et al. Blood-based profiles of DNA methylation predict the underlying distribution of cell types: a validation analysis. Epigenetics 8, 816–826 (2013).

32. Quackenbush, J. Microarray data normalization and transformation. Nat. Genet. 32, 496–501 (2002).

33. Lin, C.Y. et al. Transcriptional amplification in tumor cells with elevated c-Myc. Cell 151, 56–67 (2012).

34. Bar-Joseph, Z., Glitter, A., Simon, I. Studying and modeling dynamic biological processes using time-series gene expression data. Nat. Rev. Genet. 13, 552–564 (2012).

35. Edgar, R., Momrachev, M., Lash, A.E. Gene Expression Omnibus: NCBI gene expression and hybridization array data repository. Nucleic Acids Res 30, 207–210 (2002).

36. Gautier, L., Cope, L., Bolstad, B.M., Irizarry, R.A. Affy—analysis of Affymetrix GeneChip data at the probe level. Bioinformatics 20, 307–315 (2004).

37. Love, M.I., Huber, W., Anders, S. Moderated estimation of fold change and dispersion for RNA-Seq data with DESeq2. bioRxiv doi:10.1101/002832 (2014).

38. Frazee, A.C., Langmead, B., Leek, J.T. ReCount: a multi-experiment resource of analysis-ready RNA-seq gene count datasets. BMC Bioinformatics 12, 449 (2011).

39. Aryee, M.J. et al. Minfi: a flexible and comprehensive Bioconductor package for the analysis of Infinium DNA methylation microarrays. Bioinformatics 30, 1363–1369 (2014).

40. Hekstra, D., Taussing, A.R., Magnasco, M., Naef, F. Absolute mRNA concentrations from sequence-specific calibration of oligonucleotide arrays. Nucleic Acids Res 31, 1962–8 (2003).

